# Retinal waves align the concentric orientation map in mouse superior colliculus to the center of vision

**DOI:** 10.1101/2022.01.26.477810

**Authors:** Kai Lun Teh, Jérémie Sibille, Jens Kremkow

## Abstract

Neurons in the mouse superior colliculus (SC) are arranged in an orientation preference map that has a concentric organization, which is aligned to the center of vision and the optic flow experienced by the mouse. The developmental mechanisms that underlie this functional map remain unclear. Here, we propose that the spatiotemporal properties of spontaneous retinal waves during development provide a scaffold to establish the concentric orientation map in the mouse SC and its alignment to the optic flow. We test this hypothesis by modelling the orientation-tuned SC neurons that receive ON/OFF retinal inputs. Our results suggest that the stage III retinal wave properties, namely OFF delayed response and the wave propagation direction bias, are key factors that regulate the spatial organization of the SC orientation map. Specifically, the OFF delay mediates the establishment of orientation-tuned SC neurons by segregating their ON/OFF receptive subfields, the wave-like activities facilitate the formation of a concentric pattern, and the wave direction biases align the orientation map to the center of vision. Taken together, our model suggests that retinal waves may play an instructive role in establishing functional properties of SC neurons and provides a promising mechanism for explaining the correlations between the optic flow and the SC orientation map.

## INTRODUCTION

The superior colliculus (SC) is a midbrain structure that is important for visually guided behaviors. The neurons in the mouse SC are selective for stimulus attributes like orientation, direction, and ON/OFF polarity (Barchini et al., 2018; Basso et al., 2021; Gale and Murphy, 2014, 2018; Inayat et al., 2015; Lee et al., 2020; Wang et al., 2010). The representations of these stimulus features are arranged in functional maps, such as the orientation preference map (OPM; Ahmadlou and Heimel, 2015; Feinberg and Meister, 2015) and direction preference map (Li et al., 2020; de Malmazet et al., 2018; but see Chen et al., 2021). Interestingly, the SC OPM is organized in a concentric pattern centered around the nose position (Fig. 1A-B; Ahmadlou and Heimel, 2015), reminiscent of the optic flow experienced by the animals (Ge et al., 2021). However, the mechanisms that give rise to the concentric OPM in the SC remain elusive.

**Figure 1:**
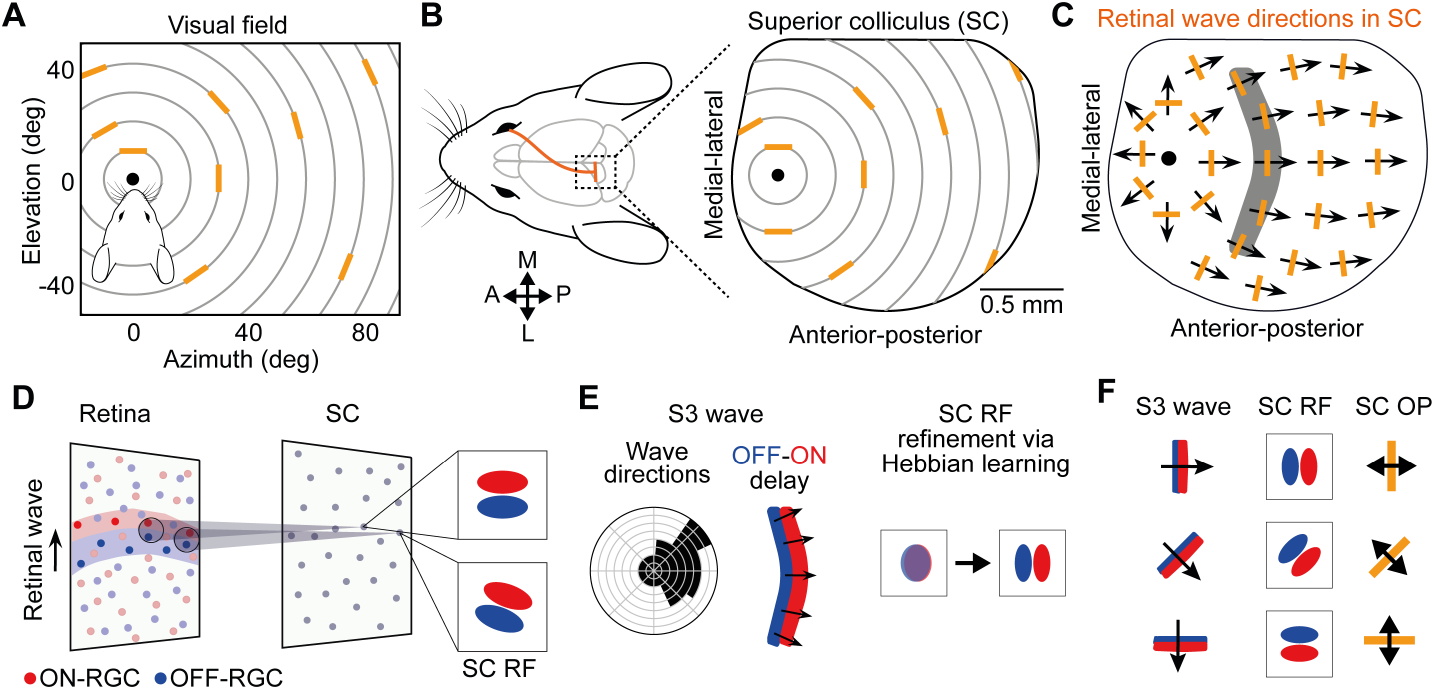
Hypothesis and model. (A-C) Hypothesis: Concentric orientation preference map (OPM) is shaped by the retinal waves during development. (A) Orientation preference of SC neurons (orange bars) plotted in different parts of the visual field. The orientation preferences in the SC are organized in a concentric pattern around the center of vision (nose position). Gray circles indicate the concentric angles, which are orthogonal to the radial angles, plotted continuously at a fixed distance from the center of vision (black dot). Data adapted from Ahmadlou and Heimel (2015). (B) The retinotopic mapping of the orientation preferences shown in A to the mouse SC. (C) Retinal waves during development. The radial retinal wave flow pattern with its center (black dot) located at the anterior part of the SC. Arrows represent the net wave vectors, which indicates the local wave directional bias. The local retinal wave vectors resemble the optic flow (Ge et al., 2021), but is orthogonal to the concentric orientation preferences (orange bars). (D-F) Computational model of retinocollicular circuit development. (D) ON- and OFF-RGC afferent inputs converge onto postsynaptic SC neurons. The receptive field (RF) of the SC neurons is fine-tuned by the presynaptic RGC activities. (E) Developmental mechanism underlying the ON-OFF segregation. During S3 waves, the wave propagation direction bias and the delayed OFF-RGC responses result in the net co-activation of the ON and OFF RGCs in the area covered by the waves, with the ON wave leading the OFF wave (left). This co-activation of the ON and OFF RGCs forms the basis for strengthening the synaptic connections between the RGCs and SC neurons using the Hebbian learning mechanism, thus facilitating the emergence of orientation-tuned RF (right). (F) The orientation preference (OP) of the SC neuron (right) depends on the configuration of the ON and OFF receptive subfields (middle), which in turn depends on the net local bias of the retinal wave direction (left).

The main afferent input to the superficial SC arises from retinal ganglion cells (RGC), the output neurons of the retina. There is growing evidence that the topographic organization in the SC emerged during the developmental period (Gribizis et al., 2019; Ge et al., 2021). During development of the visual system, there are spontaneously evoked activities, termed retinal waves, spreading across the developing retinas. The retinal waves have been shown to play important roles in refining the neural circuits of the visual systems (Huberman et al., 2008a). Specifically, the stage II (S2) retinal waves are responsible for eye-specific segregation and gross retinotopic refinement (Chandrasekaran et al., 2005; Arroyo and Feller, 2016) whereas the stage III (S3) retinal waves are known for ON-OFF segregation in the retinas and fine-scale retinotopic refinement (Chen and Regehr, 2000; Hooks and Chen, 2006; Jaubert-Miazza et al., 2005). However, it is largely unknown whether retinal waves also play a role in establishing the functional maps in the SC.

During development, retinal waves strongly drive the SC activity (Ackman et al., 2012; Gribizis et al., 2019). Interestingly, the retinal waves do not propagate in a random manner across the retina, but show a clear bias in their propagation directions. More specifically, the waves tend to propagate in the temporal-to-nasal directions in the retina, corresponding to the anterior-to-posterior directions in the SC (Ackman et al., 2012; Gribizis et al., 2019; Ge et al., 2021). When mapped onto the SC surface, the net wave flow pattern, which indicates the net local bias of the wave propagation directions, resembles the optic flow experienced by the animals (Ge et al., 2021) and matches the concentric OPM observed *in vivo* (Fig. 1C; Ahmadlou and Heimel, 2015). While both S2 and S3 waves show direction biases to some extent, only S3 waves have distinct ON/OFF responses, where the ON and OFF RGCs are activated asynchronously, with OFF RGCs having a delay relative to ON RGCs (Kerschensteiner and Wong, 2008). This asynchronous ON/OFF activation results in the co-activation of ON and OFF RGCs from neighboring areas, providing a plausible mechanism for establishing orientation-tuned receptive field (RF) in the postsynaptic SC neurons. Therefore, we hypothesize that the concentric pattern is globally coordinated by the wave propagation direction biases and the orientation preference of the SC neurons is locally established by the OFF delay. That is, we propose that the S3 retinal waves play a role in shaping the SC OPM during development. Our hypothesis predicts that 1) the OFF delay plays an important role in shaping the RF of the postsynaptic SC neurons and 2) the wave propagation direction biases provide a scaffold for the formation of functional maps.

In this work, we test this hypothesis using a computational model (Fig. 1D-F) and systematically analyze the effects of these two S3 wave properties. With our model, we show that a proper OFF delay is needed for creating well-tuned RFs. We also found that wave-like activities naturally give rise to a radial wave flow pattern, which in turn facilitates the establishment of a concentric OPM. Finally, our model discovers a surprising role of the wave propagation direction biases in aligning the OPM to the concentric angles of the visual field centered at the putative nose position (center of vision: 0° azimuth and 0° elevation), providing a promising mechanism for explaining the observations by Ahmadlou and Heimel (2015). Together, our model outlines the interplay between seemingly unrelated developmental mechanisms in shaping the SC OPM, thus provides a direct mapping from developmental features to functions in the SC. In other words, our findings support the notion that the retinal waves instruct the formation of the functional circuits in the SC.

## RESULTS

The SC neurons receive afferent inputs from a variety of RGC types. Depending on the functional type, the RGC axonal arbors organize in different SC layers (Dhande and Huberman, 2014). Among the RGC types that project to SC are the classical ON-center and OFF-center RGCs that are not orientation-tuned, and which drive ON and OFF responses in SC neurons (Wang et al., 2010). Here we model the convergence of these ON/OFF RGC inputs for establishing the orientation preference of SC neurons (Fig. 1D), similar to how the convergence of ON/OFF inputs established tuning properties in other parts of the visual system (Reid and Alonso, 1995; Paik and Ringach, 2011; Kremkow et al., 2016). This assumption is in line with results showing that while direction-selective neurons are concentrated in the superficial layers (Inayat et al., 2015; Ito et al., 2017), orientation-selective neurons are more evenly distributed across all SC layers (Ito et al., 2017), with the ON/OFF axons preferentially terminating in the lower layers of the superficial SC (Hong et al., 2011; Huberman et al., 2008b). We further assume that the spatial organization of the RFs is linked to the statistics of the retinal waves during development such that retinal waves shape the segregation of ON/OFF inputs in a refinement process. This assumption is supported by results showing that retinal waves are important for the refinement of the RGC axons during development (Chandrasekaran et al., 2005; Cheng et al., 2010; Huberman et al., 2008b).

Subsequently, we ask what could be the underlying mechanisms that refine the RF of SC neurons and establish orientation preference during development? It has been reported that the ON- and OFF-RGC activities during S3 waves are temporally anti-correlated where OFF RGCs are active with a delay of ~ 1 s relative to the ON RGCs (Fig. 1E; Kerschensteiner and Wong, 2008). This asynchronous pattern of ON/OFF RGC activation has been proposed to play a role in ON-OFF segregation in the lateral geniculate nucleus (Kerschensteiner and Wong, 2008; Gjorgjieva et al., 2009). However, its role in the SC is still unclear. Here we propose that this OFF delay plays an instructive role in establishing SC orientation preferences by providing a spatial template that selectively strengthens the ON inputs from the leading ON-wave region and OFF inputs from the following OFF-wave region with little overlap, thereby segregating the ON and OFF subfields of the SC neurons (Fig. 1D). Hebbian learning with subtractive normalization (Erwin and Miller, 1998; Lee et al., 2002; Miller, 1994; Miller and MacKay, 1994) was implemented for strengthening the co-activated presynaptic RGC inputs to the postsynaptic SC neurons (Fig. 1D-E). In this framework, the OFF delay together with the bias in wave propagation direction establish the orientation preference by fine-tuning the spatial organization of the ON/OFF subfields (Fig. 1F).

In summary, the model links the retinal wave propagation and OFF delay via Hebbian learning to the RF structure of the SC neurons. In the following, we will first demonstrate how the RF structure of local single SC neurons is dependent on the OFF delay and the local wave propagation direction bias. After that, we will show how these wave features give rise to a smooth SC OPM and how their global statistics affect the map properties.

### OFF delay and local wave directional bias establish well-tuned SC neurons

To start examining the role of the net local retinal wave vector in establishing the RF properties of the postsynaptic SC neurons, we systematically varied the OFF delay (*τ*_OFF_) from 0 s to 3.5 s and the retinal wave propagation direction bias from 0 (no bias, all directions equally probable) to 1 (only waves with the same direction) (Fig. 2A). Retinal waves propagating from eight equally spaced directions towards the center with biases in the range [0, 1] were implemented to establish the RF of the central SC neuron. The activity of the RGCs was simulated with a response kernel with a constant firing rate (Eq. 9). For overlapping ON/OFF waves (*τ*_OFF_ = 0 s), the SC RF failed to segregate into ON and OFF subfields, even with high wave propagation direction biases (Fig. 2A-B). This could be explained by the Hebbian mechanism as under this configuration both ON and OFF inputs at the same spatial location will increase their strength together and therefore cancel out each other when converging on the SC neuron. The segregation of the ON and OFF subfields, which is established by the spatial segregation of the ON/OFF inputs, emerged once there is an OFF delay (Fig. 2B). The optimal OFF delay is around 1-2 s, which is consistent with the experimental observations (Kerschensteiner and Wong, 2008). Thus, the OFF delay during the S3 waves is essential for segregating the ON/OFF subfields of SC neurons during development. However, what controls the spatial configuration of the ON/OFF subfields? When the local retinal wave directions are uniformly distributed, the SC RF failed to develop strong ON/OFF subfields even with optimal OFF delays (Fig. 2A, first column of RFs). Increasing the local wave directional bias increased the SC RF contrast and is maximum for an OFF delay of around 1-2 s and for retinal waves with strong directional bias.

**Figure 2:**
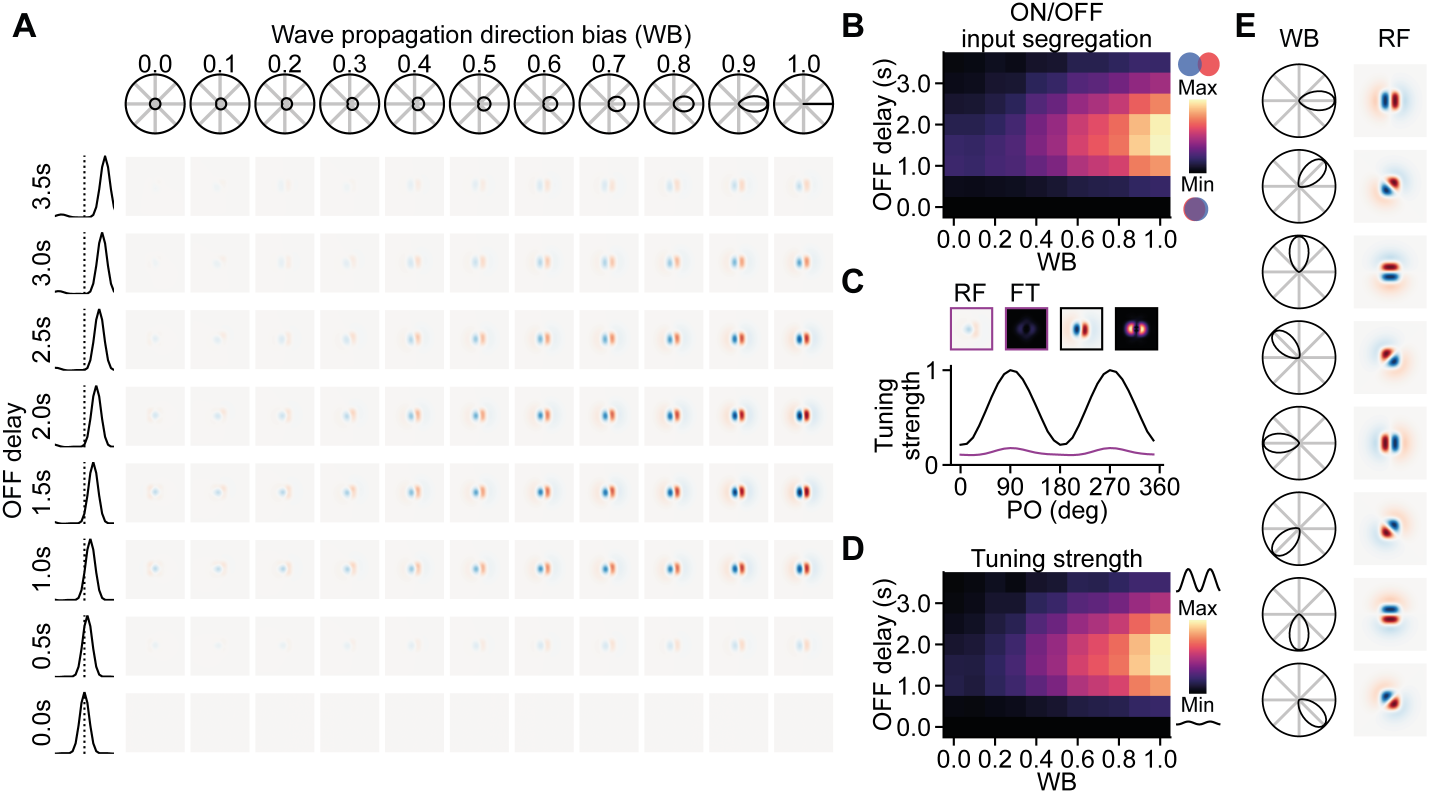
ON/OFF retinal waves shape the receptive field properties of SC neurons. (A) The effects of OFF delay and net local wave directional bias on the RF. The OFF delay of the S3 waves is needed for the separation of the ON and OFF receptive subfields. As long as there is OFF delay, higher local wave directional bias gives more prominent RF, with the OFF delays of 1.5 s and 2 s give the most prominent RF. Leftmost column: the OFF-to-ON cross-correlogram for 0 s (bottom) to 3.5 s OFF delays (top). Topmost row: the polar probability distribution of the wave directions corresponding to the bias indices. (B) The corresponding ON-OFF input segregation for the RFs in A. (C) The tuning curves of two RFs with wave propagation direction biases of 0.1 (purple) and 1 (black), both with OFF delay of 1.5 s. Insets are the corresponding RFs and their Fourier transform (FT). (D) The corresponding tuning strengths for the RFs in A. (E) The net local wave directional bias determines the final configuration of the ON and OFF receptive subfields, where the ON subfield is leading the OFF subfield in the direction of the net wave vector.

The ON-OFF segregation and orientation tuning strength of the RF have the same trend where the segregation and the tuning strength increase as the wave directional bias increases and the optimal OFF delay occurred at around 1-2 s (Fig. 2B, D). The optimal OFF delay for the RF contrast, ON-OFF segregation, and tuning strength is dependent on the size of the integration site (Eq. 14), where smaller integration site will result in shorter optimal OFF delay (data not shown). Figure 2C compares two tuning curves obtained from the Fourier transform of the corresponding RFs (Eq. 29) established with different wave directional biases.

To test whether our model could generate orientation-tuned SC neurons to any direction of retinal waves, we simulated the retinal waves with bias to eight different directions. As expected, the SC ON/OFF subfields arranged to the direction of the net wave vector with the ON subfield leading the OFF subfield (Fig. 2E). This shows that the OFF delay will result in the co-activation of neighboring ON and OFF RGC inputs. Therefore, with the Hebbian mechanism, the OFF delay helped segregate and establish the ON and OFF receptive subfields of the postsynaptic SC neurons, which in turn determined the resulting orientation preference of those SC neurons (Fig. 2E, right). Thus, this proposed model could establish orientation-tuned RFs in the postsynaptic SC neuron with spatial configuration shaped by the direction of the net local wave vector.

Taken together, biological realistic values of the OFF delay (1-2 s) observed during S3 retinal waves with sufficient local wave directional bias can establish a well-tuned RF in the SC neuron, with the spatial organization of the RF arranged in the biased wave direction.

### Radial wave flow pattern gives rise to concentric map organization

We propose that the OPM in the SC is determined by the pattern of the net retinal wave flow. This hypothesis is supported by the observations that the S2 and S3 retinal waves do not propagate uniformly to all directions (Ackman et al., 2012; Gribizis et al., 2019), and a net wave flow pattern has been measured (Ge et al., 2021). This net wave flow pattern indicates the net excess wave activities in a direction during these developmental stages, which could potentially shape the functional organization of the target structures. As shown in the previous section, these wave-directionality-induced local excessive activities could have an impact on the RGC-SC connections and give rise to an orientation-tuned SC RF. Now, we are going to explore the effects of these wave properties, namely the OFF delay and the wave propagation direction bias, on the overall organization of the SC OPM.

Non-wave-like retinal activities without consistent OFF delay (Fig. 3A_1_, left) resulted in random wave vectors (Fig. 3A_1_, right) and thus did not produce a smooth OPM (Fig. 3A_3_, left). Similarly, retinal waves with uniform propagation directions but with random ON/OFF activation also did not give rise to a smooth OPM (Fig. 3B_3_, left), despite having smooth wave vectors (Fig. 3B_1_, right). This indicates that the ON/OFF activation pattern (i.e. the OFF delay), and not just the wave-like activities, is also important in establishing a smooth OPM.

**Figure 3:**
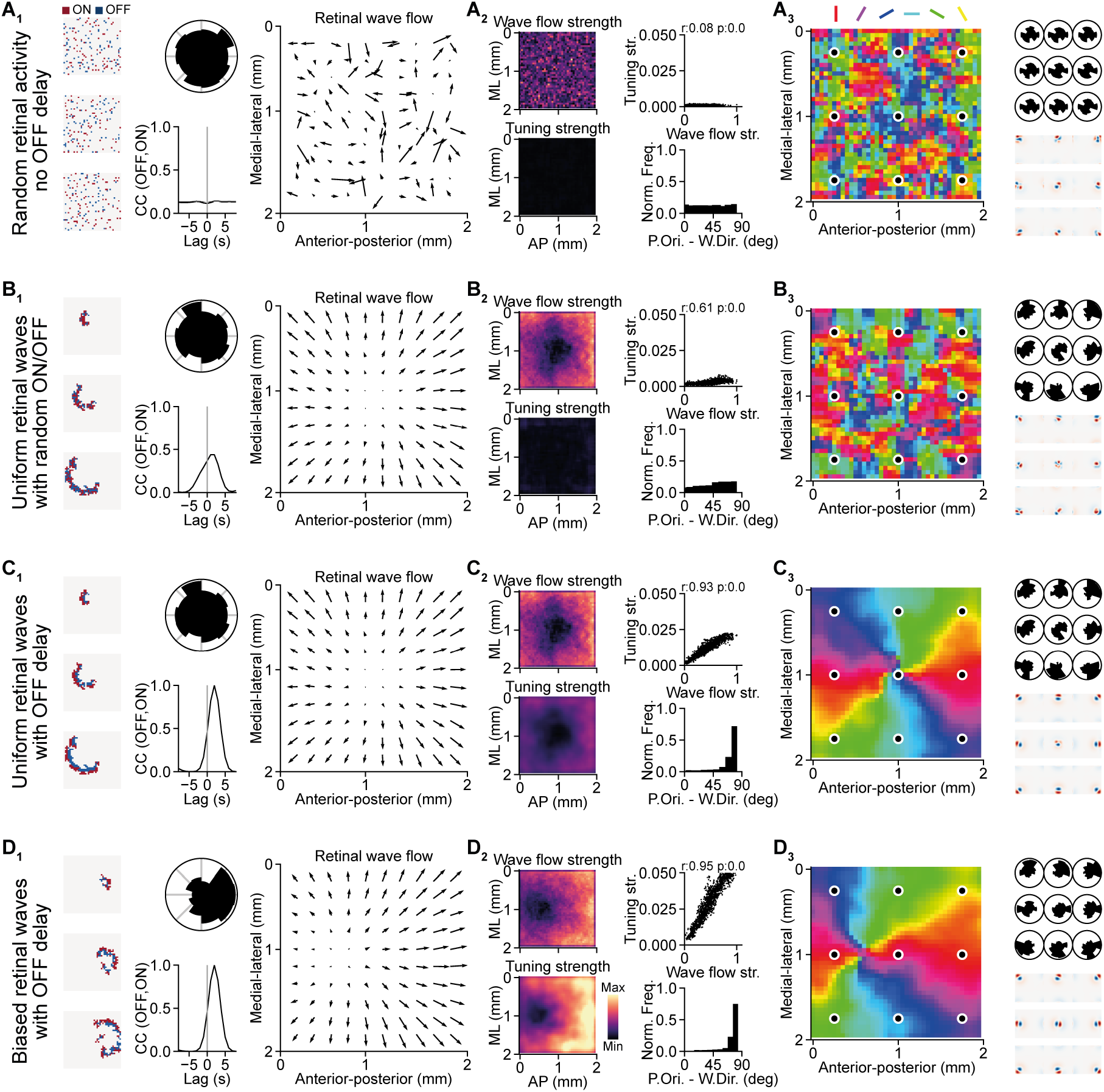
Wave-like activities with OFF delay establish a smooth SC OPM. (A) Random retinal activities. (B) Uniform retinal waves with random ON/OFF activation. (C) Uniform retinal waves with OFF delay. (D) Biased retinal waves with OFF delay. (A_1_-D_1_) Left: examples of retinal waves. Middle: distribution of wave propagation directions (top) and the cross-correlogram of the OFF relative to ON RGC activities (bottom). Right: the retinal wave flow. (A_2_-D_2_) Left: the strength of the retinal wave flow (top) that reflects the local bias of wave propagation directions and the orientation tuning strength (bottom). Right: the relationship between the wave flow strength and the orientation tuning strength (top) and the distribution of degree difference between the orientation preference and the wave propagation direction (bottom). Note that the retinal waves with OFF delay give rise to orthogonality between the orientation preference and the wave propagation direction (C_2_ and D_2_). (A_3_-D_3_) Left: the resulting OPM. Right: distribution of local wave propagation directions (top) and RFs (bottom) corresponding to the nine locations marked in the OPM.

Next, we tested the model with uniform retinal waves (Fig. 3C_1_, top middle) with *τ*_OFF_ = 1 s to see its effects. To our surprise, these retinal waves without propagation direction biases already gave rise to a smooth concentric OPM with its singularity located at the center of the model (Fig. 3C_3_, left), matching the center of the radial wave flow pattern (Fig. 3C_1_, right). A possible explanation is that although there was no global wave propagation direction bias (Fig. 3C_1_, top middle), there were still local biases at every retinotopic location (Fig. 3C_3_, top right), which formed a radial wave flow pattern (Fig. 3C_1_, right). Finally, the retinal waves with biased propagation in the anterior-to-posterior directions (which is temporal-to-nasal in the retina) and *τ*_OFF_ = 1 s (Fig. 3D_1_, top middle) produced smooth wave vectors that formed a radial pattern (Fig. 3D_1_, right) and smooth OPM with its singularity shifted towards the anterior SC (Fig. 3D_3_, left). This result is consistent with the concentric OPM observed *in vivo* (Ahmadlou and Heimel, 2015).

Therefore, our results indicate that the wave-like retinal activities can produce wave vectors that are changing gradually across neighboring retinotopic locations and have strong magnitude (Fig. 3B_2_-D_2_, top left). With proper ON-OFF temporal structure, these wave vectors that represent local wave directional bias can give rise to RFs with strong tuning strengths (Fig. 3C_2_-D_2_, bottom left) and thus producing a smooth OPM. The resulting orientation preferences are arranged orthogonally to the directions of these net local wave vectors (Fig. 3C_2_-D_2_, bottom right). In other words, the net wave flow that has a radial pattern determines the globally concentric organization of the OPM.

### Globally biased retinal waves align the concentric map to the visual field center

In the previous section, we showed that the wave propagation direction biases could affect the position of the singularity in the concentric OPM, but what is the underlying mechanism? In our model, retinal waves propagated away from a source of asymmetric inhibition, which is inspired by the asymmetric inhibition observed in vivo (Ge et al., 2021). For each wave initiation, the source of asymmetric inhibition was sampled from a 2-D Gaussian distribution centered at the center of vision (predefined nose position). The wave directional bias decreases as the spread of the asymmetric inhibition distribution increases (Fig. 4E). Thus, the location and spread of the asymmetric inhibition could play a key role in shifting the map singularity.

**Figure 4:**
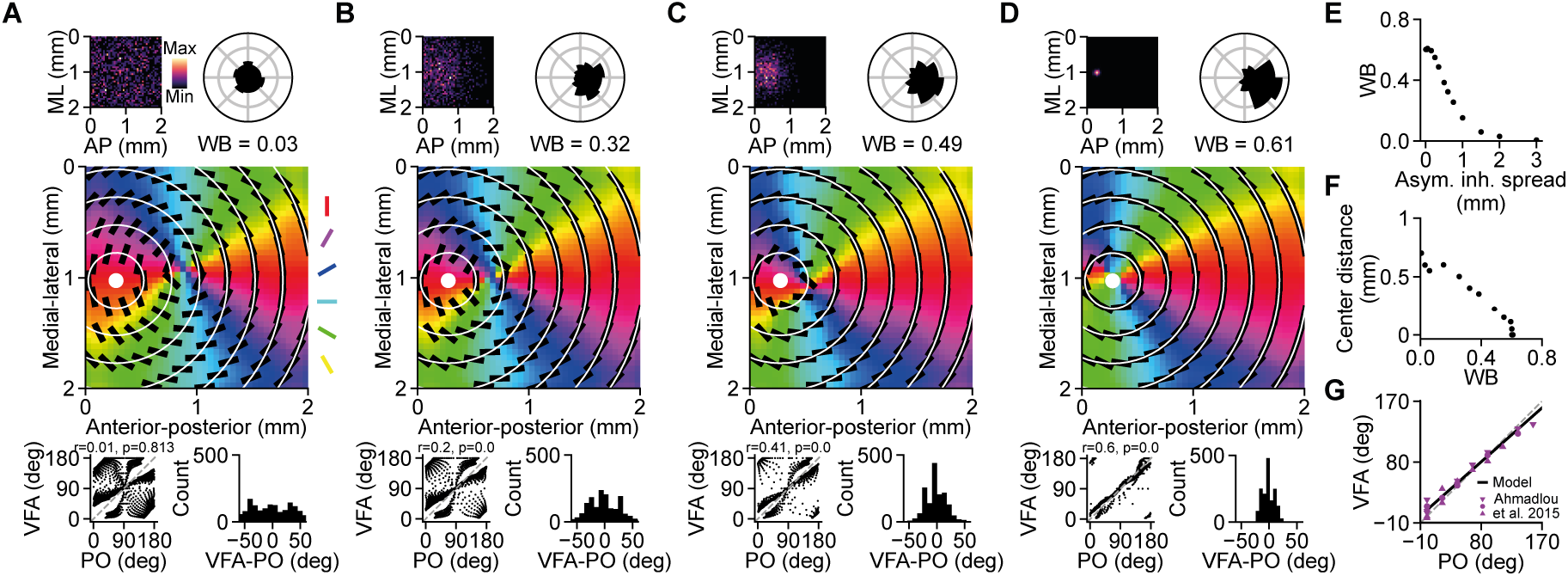
Small asymmetric inhibition spread creates a bias in the retinal waves that aligns the OPM to the concentric angles of the visual field. (A-D) Top: the distribution of the source positions of asymmetric inhibition (left) and the wave direction distribution (right). Middle: overlay of concentric angles of the visual field (circular lines) on the OPM. With higher wave direction biases, the local orientation preferences (black bars) are more aligned to the concentric visual field angles. The white dot indicates the visual field center. Bottom: the concentric visual field angles against the orientation preference (left) and the frequency distribution of their angular differences (right). (E) The relationship between the wave propagation direction biases and the spreads of the asymmetric inhibition distribution. Smaller asymmetric inhibition spreads give higher bias in the wave propagation directions. (F) The distance between the visual field center and the OPM singularity decreases as the wave direction bias increases. (G) The model with wave direction bias of 0.61 is consistent with the in vivo observation. Data replotted from Ahmadlou and Heimel (2015).

To systematically study the role of the asymmetric inhibition on the location of the OPM singularity, we gradually decreased the spread of the asymmetric inhibition distribution (Fig. 4A-D, top left). As expected, the location of the singularity gradually changed from the center of the SC towards the center of the asymmetric inhibition distribution. It is worth noting that shifting the center of an asymmetric inhibition distribution with a small spread could also shift the singularity of the resulting OPM (data not shown). However, instead of changing the center position of the asymmetric inhibition distribution, we fixed its center at the center of vision and changed its spread from 2000 *μ*m (Fig. 4A), which is equivalent to a uniform distribution that has no overall asymmetric inhibition, to 50 *μ*m (Fig. 4D), which gives high asymmetric inhibition originated from the visual field center. Nevertheless, in both cases the highly localized sources of asymmetric inhibition around the visual field center is the key factor that gives rise to wave direction biases and thus shifts the OPM singularity towards the visual field center (Fig. 4D, F). Misalignment between the OPM and the visual field will result in a large angular difference between the orientation preferences and the concentric angles of the visual field (Fig. 4A-B, bottom). The waves with direction bias of around 0.61 resulted in an angular difference distribution similar to in vivo observation (Fig. 4D; Ahmadlou and Heimel, 2015). Figure 4G shows the comparison between the model and the data from Ahmadlou and Heimel (2015) on the concentric visual field angle against the preferred orientation.

### Robustness of the concentric map pattern to the activity noise

Is the proposed model sufficiently robust to background activity noise? To answer this question, we tested the model with different fractions of signal (the waves) and noise (the random background activities), which are determined by *r* and *γ* in Eq. 9 respectively, where *r* + *γ* = 1 and *r*, *γ* ≥ 0. With increasing fraction of noise, the waves became less and less prominent (Fig. 5A). As expected, the resulting OPM became less smooth with increasing noise level (Fig. 5B). An interesting observation is that the area around the singularity of the OPM got affected first by the noise, which could be explained by the relatively low wave flow strength around the map singularity (Fig. 3C_2_-D_2_).

**Figure 5:**
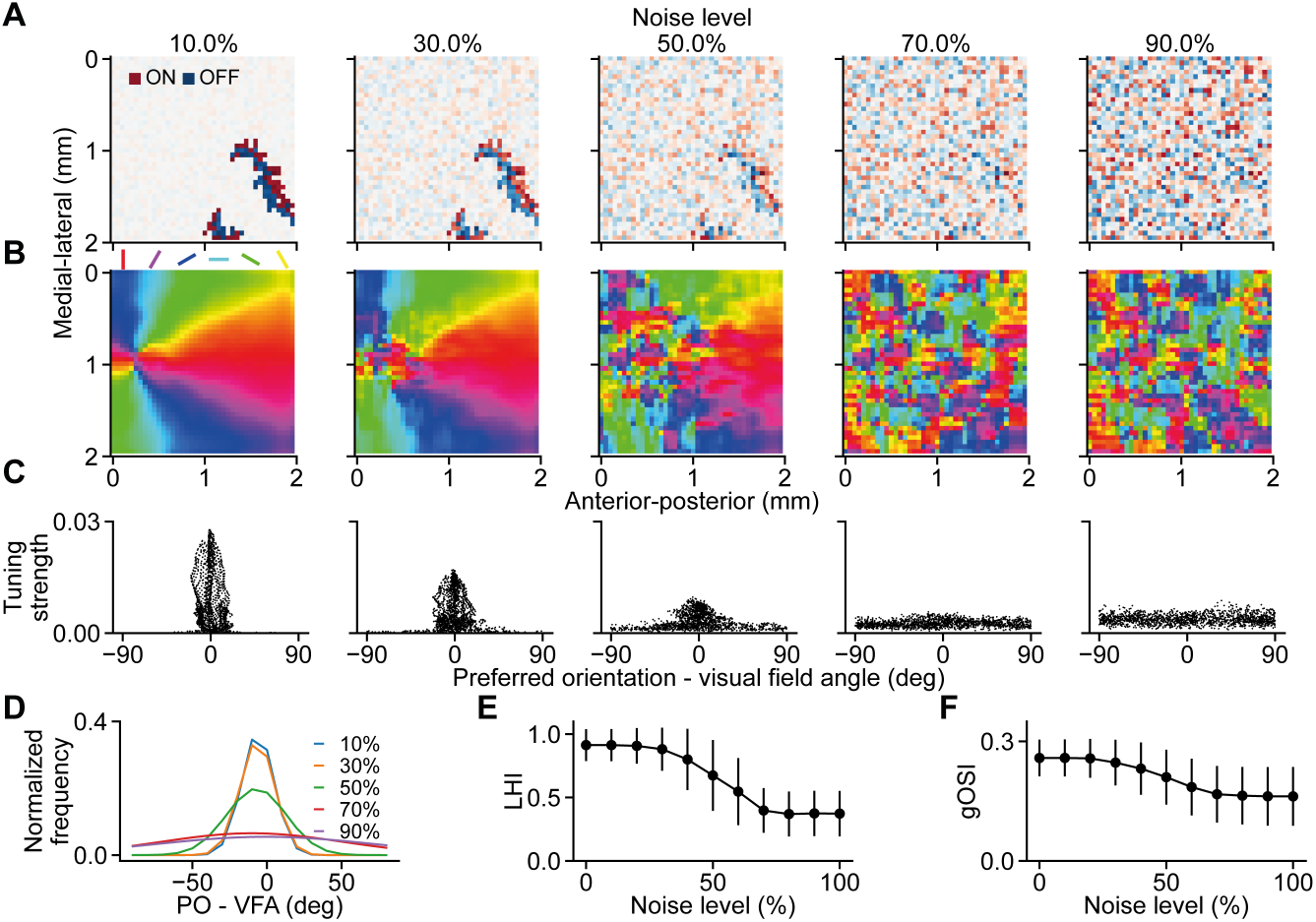
Robustness of the model to activity noise. (A) Five examples of wave frames with different levels of activity noise. (B) The resulting OPMs become less and less smooth as the activity noise increases. Note that the OPM singularity gets affected first as the noise level increases. (C) The tuning strengths vs. the angular difference between the preferred orientation and the concentric visual field angle. As the noise level increases, the tuning strength decreases and the angular difference becomes larger. (D) The distribution of the angular difference between the preferred orientation and the concentric visual field angles remained unchanged up to 30% activity noise level. (E) The local homogeneity index (LHI) decreases as the noise level increases. (F) The global orientation selectivity index (gOSI) decreases slightly as the noise level increases. Error bars indicate the standard deviation.

Additionally, as the noise level increases, the orientation tuning strength decreased, and the angular difference between the preferred orientation and the concentric visual field angle became larger (Fig. 5C). Up to 30% noise level, the angular difference remained largely intact (Fig. 5D) despite the decrease of the overall orientation tuning strength (Fig. 5C).

To quantify the effects of the noise on the map properties, we computed the local homogeneity index (LHI; Nauhaus et al., 2008) and the global orientation selectivity index (gOSI). As shown, the model is robust up to 30% activity noise level with little effects on the LHI and gOSI (Fig. 5E-F). Above 30% noise level, the LHI and the gOSI started to decrease steeply and reached the minimum at 70% noise level.

## DISCUSSION

How does the concentric orientation preference map arise in the mouse SC and why is it aligned to the center of vision? Here, we advance the notion that the SC OPM could be an emergent phenomenon that originates from retinal waves during development. The two key findings of our study are 1) the wave-like activities naturally give rise to a radial organization of the net wave flow field, which in turn facilitates the emergence of a concentric OPM (Fig. 3), and 2) the wave directionality shifts the center of the radial net wave flow, and therefore the singularity of the OPM, towards the center of the asymmetric inhibition sources (Fig. 4). Together, these two mechanisms provide an elegant explanation for the retinotopic-based orientation preference (Ahmadlou and Heimel, 2015; Feinberg and Meister, 2015) and globally concentric organization (Ahmadlou and Heimel, 2015) in the SC OPM. Since S3 retinal waves tend to propagate towards the posterior SC (Gribizis et al., 2019; Ge et al., 2021), which implies that the source of asymmetric inhibition is located at the anterior SC that corresponds to the nose position in the visual field, this suggests that the retinal wave directionality plays a role in aligning the functional maps to the optic flow, supporting the findings of our model. Interestingly, this alignment of the singularity of the OPM to the wave flow center at the anterior part of the SC could explain for the lack of a map singularity in the observation by Feinberg and Meister (2015) as they were measuring at the posterior SC.

A possible explanation for the emergence of a radial organization of the net wave flow is because the continuous spatiotemporal structure of a wave will activate neighboring RGCs simultaneously, resulting in smooth continuous change of the net local wave vectors in neighboring areas. Application of Gabazine in the retina during development, which interferes with the asymmetric inhibition that underlies the wave direction bias of S3 waves, affects the direction selectivity of SC neurons, but has little effect on SC neurons that are orientation-tuned (Ge et al., 2021). Our model can recapitulate this finding since a concentric OPM was still established during unbiased waves (Fig. 3, 4), just with a mismatch of the visual field center and the OPM singularity, which could be easily tested experimentally.

What could be the potential functional roles of aligning the concentric SC OPM to the center of vision? Recently, Ge et al. (2021) showed that the pattern of retinal waves during development matches the optic flow experienced by the adult animals. Thus, following this idea, a potential role of the concentric OPM could be important for the processing of the orientations of visual objects experienced during natural behaviors. However, it remains to be shown what benefit the match of the visual field center and the SC OPM singularity brings for the perception and visually guided behaviors of the animal. On the other hand, the distribution of tuning strengths on the established OPM might have more obvious behavioral implications. The tuning strengths become higher as the distance to the map singularity increases (Fig. 3C_2_-D_2_). That is, the orientation tuning strengths are the lowest at the singularity and the highest at the periphery of the OPM. This means that the SC is likely to be more sensitive to the orientation- and motion-related stimuli at the periphery of the visual field that are important for the detection of potential dangers and initiation of defensive behaviors. Further experiments have to be conducted to confirm this prediction.

The use of Hebbian learning to refine the RGC-SC connections, which requires strong co-activation between RGCs and SC neurons, is supported by the experimental work showing that retinal waves strongly drive SC neurons during S3 waves (Gribizis et al., 2019). This suggests that the Hebbian mechanisms could be crucial in forming the functional maps in the SC. Gribizis et al. (2019) also showed that the activities of the primary visual cortex (V1) do not correlate strongly with the S3 retinal waves, providing a plausible explanation for the salt-and-pepper organization in the mouse V1 that is different from the concentric SC OPM. These observations suggests that the retinocollicular and thalamocortical circuits may utilize different developmental mechanisms for establishing the functional organization. Consistent with this idea, we recently showed that RGC axons form mosaics in the SC that is isomorphic to the retinal mosaic (Sibille et al., 2021), while in contrast, thalamocortical axons seem to form clusters in the V1, both in species with and without OPM (Kremkow et al., 2016; Ringach, 2021). This could be related to how visual information is represented in these two different circuits; however, more work need to be done to better understand this.

The OFF delay in our model, inspired by the findings of Kerschensteiner and Wong (2008), allows the ON and OFF subfields to segregate and thereby establishes orientation-tuned SC neurons. This temporally delayed mechanism might play a role in establishing different functional features via strengthening different RGC types. For example, it could be that transient and sustained RGC types are organized in a spatiotemporal way such that direction selectivity can emerge from the selective strengthening of these RGC types (Lien and Scanziani, 2018). Hence, due to its simplicity, our model can be easily extended to incorporate the establishment of other functional maps, e.g. the motion direction preference map. Since the direction preference is orthogonal to the orientation preference (Li et al., 2020; but see de Malmazet et al., 2018 and Chen et al., 2021), which is similar to the angular difference between the net wave vector and the orientation preference (Fig. 3C_2_-D_2_), the extended model could postulate that the direction preference is parallel to the net wave vector.

Although the model can recapitulate several key findings about the SC OPM, e.g. the concentric organization and alignment to the optic flow, there are some limitations of the model. First, we used a simplified probabilistic, rather than detailed mechanistic, approach to model the asymmetric inhibition, which might overlook some key details of the asymmetric inhibition process. Besides, we assumed that the model is only retinotopically mapped from the retina to the SC and no other functional properties are established at the beginning of the simulations, which ignored the potential that the S2 retinal waves might have already shaped the functional maps to some extent. Nevertheless, it is still reasonable to believe that the simplified mechanisms of our model do contribute significantly to the shaping and maturation of the SC OPM.

Despite its simplicity, the model provides a set of predictions that could be tested *in vivo*. 1) Interfering with the wave direction bias, but leaving the OFF delay intact, should result in a different SC OPM. In case of uniform wave directions, it should be a concentric OPM with its singularity located at the center, instead of the anterior part, of the SC. This can be tested *in vivo* by applying Gabazine to block the GABAergic inhibition in the retina during development. 2) In our model, the OFF receptive subfield is oriented towards the center of the retinal wave flow, which is also the visual field center. This can be validated by mapping the ON and OFF subfields of orientation-selective SC neurons. 3) In our model, the tuning strength of SC neurons varies systematically across the SC OPM, with strongly tuned neurons being located in regions with large net wave vector. Our model predicts that the region around the OPM singularity has relatively low tuning strength (Fig. 3C-D) and is more susceptible to the noise (Fig. 5).

Many theories and mechanisms have been proposed for the establishment of OPM in the V1 of higher mammals (Swindale, 1992; Obermayer and Blasdel, 1993; Miller, 1994; Swindale, 1996; Erwin and Miller, 1998; Koulakov and Chklovskii, 2001; Ringach, 2004; Mariño et al., 2005; Kaschube et al., 2010; Paik and Ringach, 2011; Reichl et al., 2012; Kremkow et al., 2016; Jang et al., 2020; Najafian et al., 2022). However, the OPM in the SC is less well-studied. Due to the difference between the V1 and SC OPMs, where V1 has equal representation for all orientations in every retinotopic location (Hübener et al., 1997) whereas SC has retinotopically biased representation of the orientations (Ahmadlou and Heimel, 2015; Feinberg and Meister, 2015), the proposed mechanisms for the V1 OPM may not be suitable for explaining the SC OPM. Our model helps bridge this gap by showing the viability of retinal waves in shaping the SC OPM. It is well-known that retinal waves are key factors for refining the retinotopic map in the visual system, including SC (McLaughlin et al., 2003; Chandrasekaran et al., 2005; Huberman et al., 2008b). However, how functional response properties of SC neurons are established during development by the retinal waves remain largely unknown. Our modeling work now suggests that there is a much more direct link between the retinal wave flow statistics and the SC functional maps than previously thought. Thus, spontaneous activity during development not only refines the retinotopic map (Chandrasekaran et al., 2005), but has further potentials in establishing the functional organization in the target regions. Taken together, our model provides a plausible framework for associating developmental properties in developing animals to the functional organization in adult animals. The parsimonious explanations offered by our model outline a promising path towards understanding the design of the functional circuits by the developmental mechanisms, which has been a long-standing question in many different brain regions.

## ACKNOWLEDGEMENTS

We thank J.-M. Alonso and R. Kempter for helpful discussions during the project; Tatiana Lupashina and Carolin Gehr for comments on the manuscript; This work was supported by the DFG Emmy-Noether grant KR 4062/4–1 (J.K.).

## AUTHOR CONTRIBUTIONS

K.-L.T., J.S. and J.K. conceived and designed the study; K.-L.T. implemented and studied the model; K.-L.T. and J.K. analyzed the data; and K.-L.T. and J.K. wrote the manuscript with inputs from J.S.

## METHODS

### Two-layer retina-SC model

The model has two layers. The first layer modeled a retina that contains 1600 ON RGCs and 1600 OFF RGCs, each arranged in a 40 × 40 grid. The length of a pixel is 50 *μ*m. The second layer has 1600 SC neurons, also arranged in a 40 × 40 grid. The RGCs were connected to the SC neurons retinotopically (Eq. 14), i.e. the neighboring RGCs projected to similar retinotopic locations in the SC. The synaptic weights were initialized uniformly with strength of 0.1. The diameter of the integration site of RGC inputs, which is the sum of the average RGC axonal diameter and the average dendritic diameter of SC neurons, was set to 350 *μ*m (Ackman et al., 2012; Hong et al., 2011; Huberman et al., 2008b) and its radius *R*_radius_ = 175 μm was used for defining the arbor function (Eq. 14).

The retinal positions are denoted by Greek alphabets 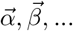 and the SC positions are denoted by Roman alphabets 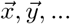, which are all 2-D vectors. Since the model assumed the presynaptic RGCs have already been retinotopically mapped to the postsynaptic SC neurons, the location vectors 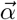 and 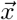 also reflect the receptive field (RF) position in the visual field 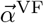 (Eq. 20). For all illustrations in this model, the top, right, bottom, and left refer to dorsal (D), temporal (T), ventral (V), and nasal (N) in the visual field, respectively. Consequently, these directions correspond to V, N, D, and T in the retina and medial (M), posterior (P), lateral (L), and anterior (A) in the SC (Fig. 1A-B; Ge et al., 2021).

Two functions that were frequently used are the 2-D Gaussian function and the polar bias index. The 2-D Gaussian function used in this paper was defined as

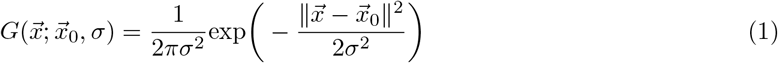

where ||.|| denotes the Euclidean norm, 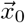 is the center, and *σ* is the spread. The polar bias index, *B*, was computed by:

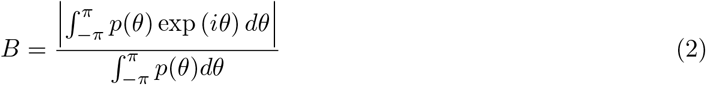

where *p* is a polar probability density function truncated for the interval (−*π*, *π*] and *p*(*θ*) is the probability for a direction *θ* ∈ (−*π*, *π*]. Since the normalization term 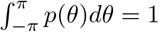, *B* is reduced to

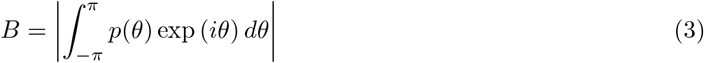

### Retinal wave model

Each wave *w* was initiated at position 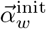 one after another by sampling from the uniformly distributed wave initiation map (Ge et al., 2021). The temporal resolution, i.e. the duration per frame of wave, was set to 0.5 s. To implement the asymmetric inhibition shown in Ge et al. (2021), a source position of the asymmetric inhibition 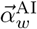 was sampled from a 2-D Gaussian distribution centered at the putative nose position (center of vision) 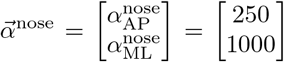 with spread *σ*^AI^ ∈ {5, 25, 50, 150, 250, 350, 500, 600, 750, 1000, 1500, 2000, 3000} *μ*m (Fig. 4).

### Wave propagation

The activation state of the RGC at the position 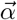 in the wave frame *t* was described by

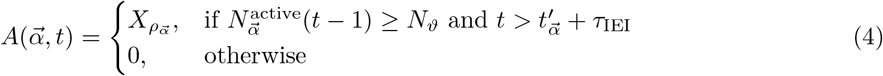

where 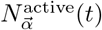 is the number of active nearest neighbors (NN) of the 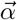 RGC at time *t*, *N_ϑ_* = 1 is the threshold number of active NNs, 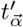 is the time of last activation for the RGC at 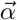, *τ*_IEI_ = 30 s is the inter-event interval (IEI; Gribizis et al., 2019), and 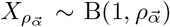 is the sample from a binomial distribution with the activation probability 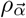.

The local area for position 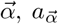, was defined as the 3 × 3 pixels centered at 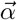. To induce directed waves, the activation probability 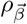 for the 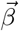 NNs of 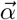 RGC was not uniformly distributed and was determined by the polar probability density function for local propagation *p*^prop^:

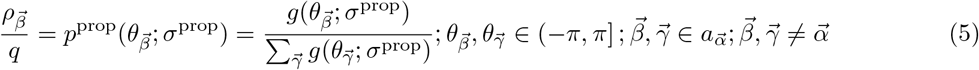

where *q* = 2.4 is a scaling factor and

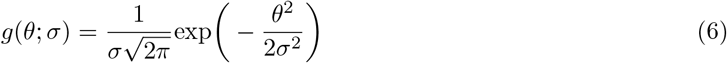

is the 1-D Gaussian function truncated for the interval (−*π*, *π*] with mean of 0 and standard deviation *σ*. The standard deviation *σ*^prop^ was numerically optimized by

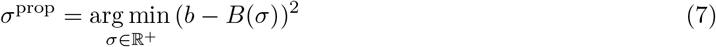

where *b* = 0.35 is the desired local propagation bias and

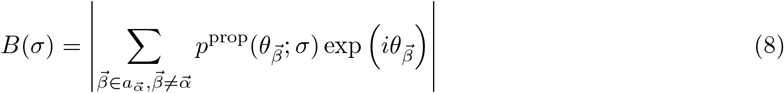

is the discrete version of Eq. 3 for calculating the bias of the local propagation probability density function *p*^prop^. 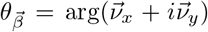, where 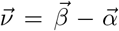. In the square grid configuration of the model, 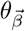 can also be expressed as *θ_k_*, where 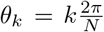 is the direction of *k*th NN at 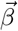 relative to the RGC at 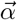 and *N* = 8 is the total number of NNs. The *b* controls the shape and size of the retinal wave, where higher *b* will produce smaller and straighter waves that resemble the S3 waves and lower *b* will produce larger, curved waves that resemble the S2 waves. The wave propagation direction was computed as 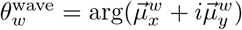, where 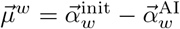. Next, the 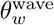 was rounded to the nearest *θ_k_* to get 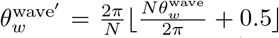. The peak probability of the *p*^prop^ was then oriented to the 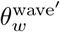 such that 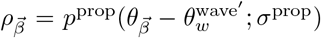. In other words, the wave *w* was propagating from its initiation position 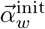 away from the source position of the asymmetric inhibition 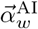.

### RGC activity

The activity of the active RGCs was modeled as firing rate represented by a response kernel defined by

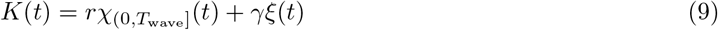

where *r* is the mean RGC firing rate, *T*_wave_ is the pixel activation duration during the wave, *γ* is the strength for noise and was set to 0 if not stated otherwise, *ξ*(*t*) is the Gaussian white noise, and *χ_T_*(*t*) is the indicator function for interval *T*:

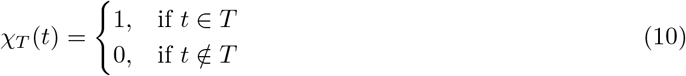

The response kernel for RGCs with polarity *P* and delay *τ_P_* was given by

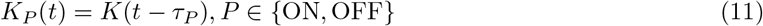

*τ*_ON_ = 0 s and *τ*_OFF_ = 1 s were used in the simulations to mimic the delayed OFF activation (Kerschensteiner and Wong, 2008) unless stated otherwise.

The RGC activity of polarity *P* at location 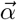 at time *t* was computed by

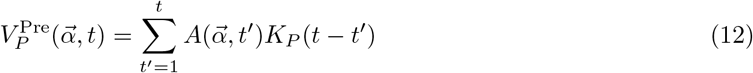

This equation implies that the activity of OFF RGCs is just a copy of the ON-RGC activity with a delay of *τ*_OFF_ – *τ*_ON_.

### Learning mechanism

Since the activity of postsynaptic SC neurons is highly correlated to the presynaptic stage III RGC activity (Gribizis et al., 2019), Hebbian learning rule (Erwin and Miller, 1998; Lee et al., 2002) with subtrative normalization (Goodhill, 1993; Goodhill and Barrow, 1994; Miller, 1994; Miller and MacKay, 1994; Elliott and Shadbolt, 2002; Elliott, 2003) was used to update the synaptic weights between the presynaptic RGCs and the postsynaptic SC neurons.

The postsynaptic activity of SC neuron at location 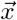 is a weighted sum of all its presynaptic inputs 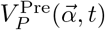:

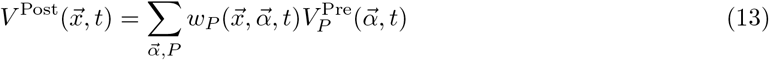

where 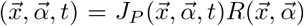 is the synaptic weight at time *t* from RGC of polarity *P* at 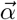 to SC neuron at 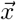. 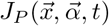 gives the synaptic weight of all presynaptic-postsynaptic pairs and

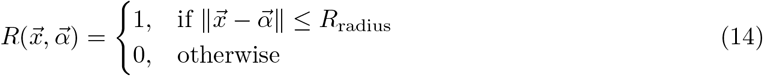

is the arbor function (input integration site) for ensuring that the influence is constrained by the retinotopy. *R*_radius_ = 175 *μ*m is two times the radius of the axonal arbor estimated from Ackman et al. (2012), Hong et al. (2011), and Huberman et al. (2008b).

The initial synaptic weight *w*_0_ was set to 0.1. The synaptic weight update was computed by

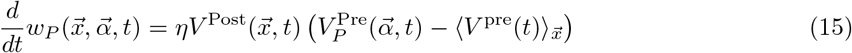

where *η* is the learning rate and

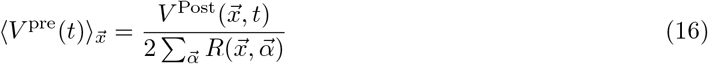

is the weighted average activity of the presynaptic RGCs that have projection to the postsynaptic SC neuron at location 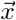. To speed up the computation, we binned the time *t* into trials *T_i_* = *iT*_trial_ and updated the synaptic weights after each trial by averaging over the presynaptic activities. By substituting Eq. 13 and Eq. 16 into Eq. 15, we have

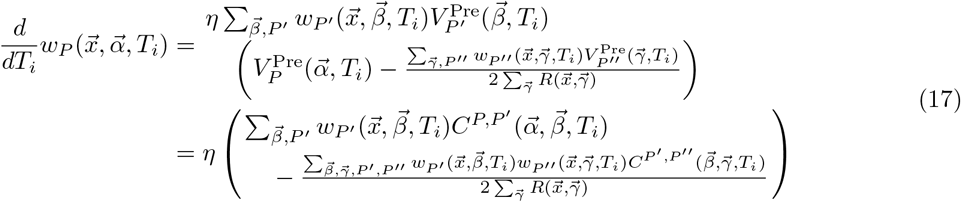

where

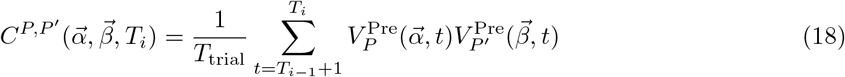

*T*_trial_ = *nt* is the trial duration and *i*, 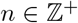. To prevent the synaptic weights from becoming negative, the synaptic weights were constrained by

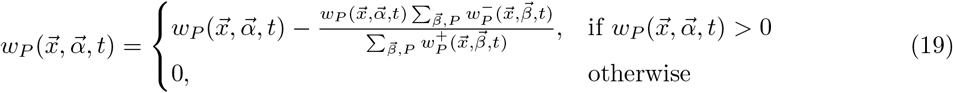

after updating the synaptic weights of each trial. The 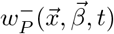 gives the absolute value of the negative weights and the 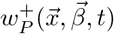 gives the positive weights from the *P* RGC at 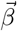 to the postsynaptic SC neuron at 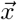. Together, the subtractive normalization (Eq. 15) and the weight constraint (Eq. 19) ensured that the total synaptic weight between all RGCs and each SC neuron is conserved, i.e. 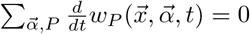, while keeping all synaptic weights non-negative. These mechanisms were used to mimic the response homeostasis (Chandrasekaran et al., 2007).

### Receptive field

The RF of RGC at location 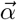 with polarity *P*, 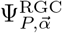, was modeled as 2-D difference of Gaussian in the visual field coordinates 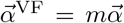 with *m* = 2.5 as the scaling factor from retinal to visual field coordinates:

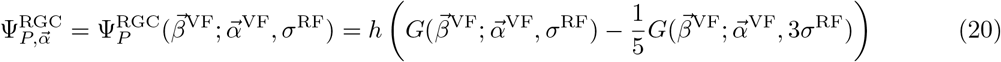

where

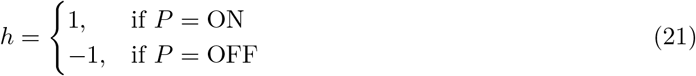

The *σ*^RF^ = 17 indicates the width of the RGC RF in visual field pixels. The scale is set to 0.6 degrees per visual field pixel.

The RF of the SC neuron at location 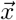 is the weighted sum of all its presynaptic RGC RFs at time *t*

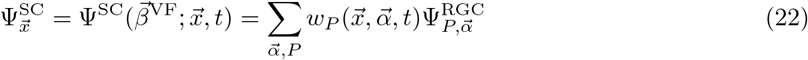

The contrast of the SC RF, 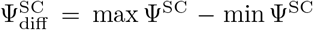, is the difference between its maximal and minimal values.

### Analyses

#### Net wave vector

The local net wave vector of location 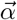 over its local area 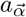 (area of 3 × 3 pixels centered at 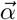) was computed by

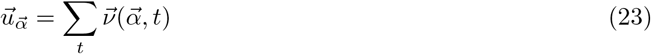

where

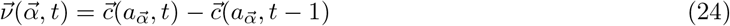

is the instantaneous wave vector at time *t* and

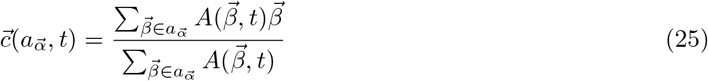

is the center of mass for the activity within the area 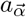 at time *t*. The magnitude of the wave vector is 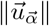 and the direction is 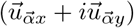.

#### Wave propagation direction bias

The bias of the wave propagation direction, *B*^wave^, was computed by:

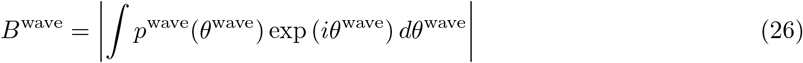

where *p*^wave^ is the normalized frequency distribution of the wave propagation directions *θ*^wave^.

#### Orientation map analyses

The orientation preference *θ*_pref_ = arg(*μ*)/2 of the SC neuron was estimated from the Fourier transform of its RF, 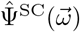 (Paik and Ringach, 2011), where

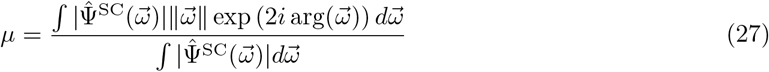

and 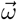 is the vector of spatial frequency components. The global orientation selectivity index (gOSI) was computed by

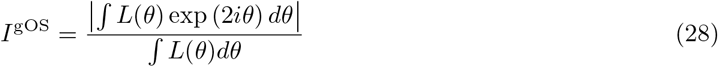

where

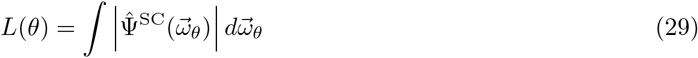

is the orientation tuning curve (Fig. 2C) obtained by summing over the frequencies 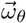 for a given orientation 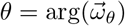. The tuning strength *L*_max_ = max *L*(*θ*) is the maximal value of the tuning curve *L*(*θ*). The singularity of the concentric OPM was estimated from the minimal wave flow magnitude.

#### ON-OFF segregation index

The segregation of the ON and OFF inputs to SC neuron at location 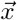 at time *t* was computed by

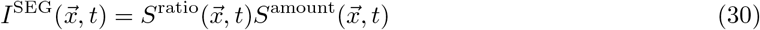

where

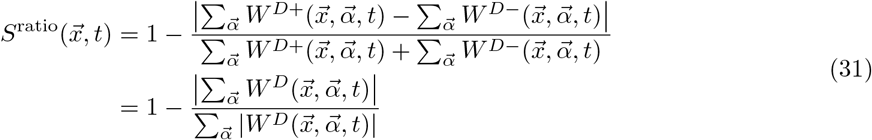

measures how balanced are the ON and OFF inputs (perfectly balanced means ON-to-OFF ratio at location 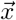 is 1 and thus 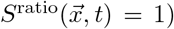), i.e. the inverse of the overall net ON or OFF input to the postsynaptic SC neuron at location 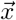 (the inverse of absolute ON-OFF dominance) and

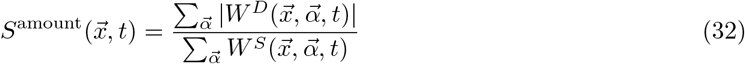

measures the amount of net ON and OFF inputs from each 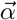 in comparison to total inputs, which is the amount of non-overlapping ON and OFF inputs to the postsynaptic SC neuron at location 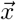. 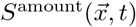 is equivalent to DSEG (degree of segregation) in Lee et al. (2002) and the non-absolute version of 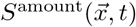 is equivalent to SIGN in Lee et al. (2002) and segregation index in Gjorgjieva et al. (2009). 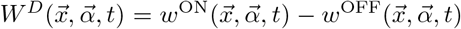 gives the net ON (positive) or OFF (negative) input from location 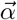 to the SC neuron at location 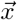 and 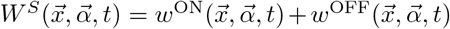 gives the total ON and OFF inputs from location 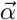 to the SC neuron at location 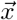. By substituting Eq. 31 and Eq. 32 into Eq. 30, we have

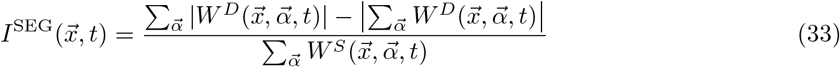

Both 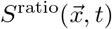 and 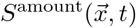 have values in range [0, 1], therefore 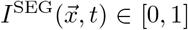. Similarly, the same formulation as Eq. 33 can be used to compute the ON-OFF subfield segregation 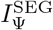 between the *P* subfields of the SC RF 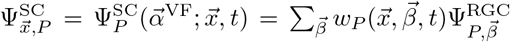 by replacing the 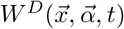 with 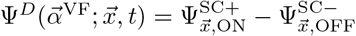 and 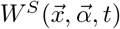 with 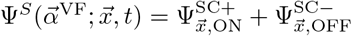:

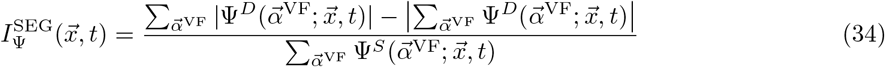

where 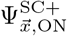 denotes the positive part of the ON subfield and 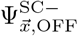 denotes the absolute value of the negative part of the OFF subfield.

#### Local homogeneity index

The local homogeneity index (LHI) is a measure of how similar the local orientation preferences are (Nauhaus et al., 2008). The LHI for location 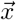 was computed by

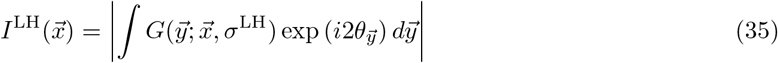

where 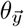 is the orientation preference at location 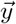 and *σ*^LH^ = 100 *μ*m.

## References

Ackman, J. B., Burbridge, T. J., and Crair, M. C. (2012). Retinal waves coordinate patterned activity throughout the developing visual system. Nature, 490:219–225.

Ahmadlou, M. and Heimel, J. A. (2015). Preference for concentric orientations in the mouse superior colliculus. Nature Communications, 6(6773):1–11.

Arroyo, David, A. and Feller, M. B. (2016). Spatiotemporal features of retinal waves instruct the wiring of the visual circuitry. Frontiers in Neural Circuits, 10(54):1–7.

Barchini, J., Shi, X., Chen, H., and Cang, J. (2018). Bidirectional encoding of motion contrast in the mouse superior colliculus. eLife, 7:226.

Basso, M. A., Bickford, M. E., and Cang, J. (2021). Unraveling circuits of visual perception and cognition through the superior colliculus. Neuron, 109(6):918–937.

Chandrasekaran, A. R., Plas, D. T., Gonzalez, E., and Crair, M. C. (2005). Evidence for an instructive role of retinal activity in retinotopic map refinement in the superior colliculus of the mouse. The Journal of Neuroscience, 25(29):6929–6938.

Chandrasekaran, A. R., Shah, R. D., and Crair, M. C. (2007). Developmental homeostasis of mouse retinocollicular synapses. The Journal of Neuroscience, 27(7):1746–1755.

Chen, C. and Regehr, W. G. (2000). Developmental remodeling of the retinogeniculate synapse. Neuron, 28(3):955–966.

Chen, H., Savier, E. L., DePiero, V. J., and Cang, J. (2021). Lack of evidence for stereotypical direction columns in the mouse superior colliculus. The Journal of Neuroscience, 41(3):461–473.

Cheng, T.-W., Liu, X.-B., Faulkner, R. L., Stephan, A. H., Barres, B. A., Huberman, A. D., and Cheng, H.-J. (2010). Emergence of lamina-specific retinal ganglion cell connectivity by axon arbor retraction and synapse elimination. The Journal of Neuroscience, 30(48):16376–16382.

de Malmazet, D., Kühn, N. K., and Farrow, K. (2018). Retinotopic separation of nasal and temporal motion selectivity in the mouse superior colliculus. Current Biology, 28(18):2961 – 2969.e4.

Dhande, O. S. and Huberman, A. D. (2014). Retinal ganglion cell maps in the brain: implications for visual processing. Current Opinion in Neurobiology, 24:133–142.

Elliott, T. (2003). An analysis of synaptic normalization in a general class of hebbian models. Neural Computation, 15(4):937–963.

Elliott, T. and Shadbolt, N. R. (2002). Multiplicative synaptic normalization and a nonlinear hebb rule underlie a neurotrophic model of competitive synaptic plasticity. Neural Computation, 14(6):1311–1322.

Erwin, E. and Miller, K. D. (1998). Correlation-based development of ocularly matched orientation and ocular dominance maps: determination of required input activities. The Journal of Neuroscience, 18(23):9870–9895.

Feinberg, E. H. and Meister, M. (2015). Orientation columns in the mouse superior colliculus. Nature, 519(7542):229–232.

Gale, S. D. and Murphy, G. J. (2014). Distinct representation and distribution of visual information by specific cell types in mouse superficial superior colliculus. The Journal of neuroscience: the official journal of the Society for Neuroscience, 34(40):13458 – 13471.

Gale, S. D. and Murphy, G. J. (2018). Distinct cell types in the superficial superior colliculus project to the dorsal lateral geniculate and lateral posterior thalamic nuclei. Journal of Neurophysiology, 120(3):1286 – 1292.

Ge, X., Zhang, K., Gribizis, A., Hamodi, A. S., Sabino, A. M., and Crair, M. C. (2021). Retinal waves prime visual motion detection by simulating future optic flow. Science, 373(6553):eabd0830.

Gjorgjieva, J., Toyoizumi, T., and Eglen, S. J. (2009). Burst-time-dependent plasticity robustly guides on/off segregation in the lateral geniculate nucleus. PLoS Computational Biology, 5(12):e1000618.

Goodhill, G. J. (1993). Topography and ocular dominance: a model exploring positive correlations. Biological Cybernetics, 69(2):109–118.

Goodhill, G. J. and Barrow, H. G. (1994). The role of weight normalization in competitive learning. Neural Computation, 6(2):255–269.

Gribizis, A., Ge, X., Daigle, T. L., Ackman, J. B., Zeng, H., Lee, D., and Crair, M. C. (2019). Visual cortex gains independence from peripheral drive before eye opening. Neuron, 104(4):711–723.

Hong, Y. K., Kim, I.-J., and Sanes, J. R. (2011). Stereotyped axonal arbors of retinal ganglion cell subsets in the mouse superior colliculus. The Journal of Comparative Neurology, 519:1691–1711.

Hooks, B. M. and Chen, C. (2006). Distinct roles for spontaneous and visual activity in remodeling of the retinogeniculate synapse. Neuron, 52(2):281–291.

Huberman, A. D., Feller, M. B., and Chapman, B. (2008a). Mechanisms underlying development of visual maps and receptive fields. Annual Review of Neuroscience, 31:479–509.

Huberman, A. D., Manu, M., Koch, S. M., Susman, M. W., Lutz, A. B., Ullian, E. M., Baccus, S. A., and Barres, B. A. (2008b). Architecture and activity-mediated refinement of axonal projections from a mosaic of genetically identified retinal ganglion cells. Neuron, 59(3):425–438.

Hübener, M., Shoham, D., Grinvald, A., and Bonhoeffer, T. (1997). Spatial relationships among three columnar systems in cat area 17. The Journal of Neuroscience, 17(23):9270–9284.

Inayat, S., Barchini, J., Chen, H., Feng, L., Liu, X., and Cang, J. (2015). Neurons in the most superficial lamina of the mouse superior colliculus are highly selective for stimulus direction. The Journal of Neuroscience, 35(20):7992–8003.

Ito, S., Feldheim, D. A., and Litke, A. M. (2017). Segregation of visual response properties in the mouse superior colliculus and their modulation during locomotion. Journal of Neuroscience, 37(35):8428–8443.

Jang, J., Song, M., and Paik, S. B. (2020). Retino-cortical mapping ratio predicts columnar and salt-and-pepper organization in mammalian visual cortex. Cell Reports, 30(10):3270–3279.

Jaubert-Miazza, L., Green, E., Lo, F.-S., Bui, K., Mills, J., and Guido, W. (2005). Structural and functional composition of the developing retinogeniculate pathway in the mouse. Visual Neuroscience, 22(5):661–676.

Kaschube, M., Schnabel, M., Löwel, S., Coppola, D. M., White, L. E., and Wolf, F. (2010). Universality in the evolution of orientation columns in the visual cortex. Science, 330:1113–1116.

Kerschensteiner, D. and Wong, R. O. L. (2008). A precisely timed asynchronous pattern of on and off retinal ganglion cell activity during propagation of retinal waves. Neuron, 58(6):851–858.

Koulakov, A. A. and Chklovskii, D. B. (2001). Orientation preference patterns in mammalian visual cortex: A wire length minimization approach. Neuron, 29(2):519–527.

Kremkow, J., Jin, J., Wang, Y., and Alonso, J. M. (2016). Principles underlying sensory map topography in primary visual cortex. Nature, 533:52–57.

Lee, C. W., Eglen, S. J., and Wong, R. O. L. (2002). Segregation of on and off retinogeniculate connectivity directed by patterned spontaneous activity. Journal of Neurophysiology, 88:2311–2321.

Lee, K. H., Tran, A., Turan, Z., and Meister, M. (2020). The sifting of visual information in the superior colliculus. eLife, 9:e50678.

Li, Y., Turan, Z., and Meister, M. (2020). Functional architecture of motion direction in the mouse superior colliculus. Current Biology, 30:1–12.

Lien, A. D. and Scanziani, M. (2018). Cortical direction selectivity emerges at convergence of thalamic synapses. Nature, 558(7708):80–86.

Mariño, J., Schummers, J., Lyon, D. C., Schwabe, L., Beck, O., Wiesing, P., Obermayer, K., and Sur, M. (2005). Invariant computations in local cortical networks with balanced excitation and inhibition. Nature Neuroscience, 8(2):194–201.

McLaughlin, T., Torborg, C. L., Feller, M. B., and D.M., O. D. (2003). Retinotopic map refinement requires spontaneous retinal waves during a brief critical period of development. Neuron, 40(6):1147–1160.

Miller, K. D. (1994). A model for the development of simple cell receptive fields and the ordered arrangement of orientation columns through activity-dependent competition between on-and off-center inputs. The Journal of Neuroscience, 14(1):409–441.

Miller, K. D. and MacKay, D. J. C. (1994). The role of constraints in hebbian learning. Neural Computation, 6:100–126.

Najafian, S., Koch, E., Teh, K.-L., Jin, J., Rahimi, H., Zaidi, Q., Kremkow, J., and Alonso, J.-M. (2022). A theory of cortical map formation in the visual brain. bioRxiv, pages 1–71.

Nauhaus, I., Benucci, A., Carandini, M., and Ringach, D. L. (2008). Neuronal selectivity and local map structure in visual cortex. Neuron, 57:673–679.

Obermayer, K. and Blasdel, G. G. (1993). Geometry of orientation and ocular dominance columns in monkey striate cortex. The Journal of Neuroscience, 13(10):4114–4129.

Paik, S.-B. and Ringach, D. L. (2011). Retinal origin of orientation maps in visual cortex. Nature Neuroscience, 14(7):919–925.

Reichl, L., Heide, D., Löwel, S., Crowley, J. C., Kaschube, M., and Wolf, F. (2012). Coordinated optimization of visual cortical maps (i) symmetry-based analysis. PLoS Computational Biology, 8(11):e1002466.

Reid, R. C. and Alonso, J.-M. (1995). Specificity of monosynaptic connections from thalamus to visual cortex. Nature, 378:281–284.

Ringach, D. L. (2004). Haphazard wiring of simple receptive fields and orientation columns in visual cortex. Journal of Neurophysiology, 92:468–476.

Ringach, D. L. (2021). Sparse thalamocortical convergence. Current Biology, 31:1–4.

Sibille, J., Gehr, C., Benichov, J. I., Balasubramanian, H., Teh, K. L., Lupashina, T., Vallentin, D., and Kremkow, J. (2021). Strong and specific connections between retinal axon mosaics and midbrain neurons revealed by large scale paired recordings. bioRxiv, pages 1–30.

Swindale, N. V. (1992). A model for the coordinated development of columnar systems in primate striate cortex. Biological Cybernetics, 66:217–230.

Swindale, N. V. (1996). The development of topography in the visual cortex: a review of models. Network: Computation in Neural Systems, 7:161–247.

Wang, L., Sarnaik, R., Rangarajan, K., Liu, X., and Cang, J. (2010). Visual receptive field properties of neurons in the superficial superior colliculus of the mouse. The Journal of Neuroscience, 30(49):16573–16584.

